# Genome-wide analysis of anterior-posterior mRNA regionalization in *Stentor coeruleus* reveals a role for the microtubule cytoskeleton

**DOI:** 10.1101/2023.01.09.523364

**Authors:** Ashley R. Albright, Connie Yan, David Angeles-Albores, Yina Hudnall, Tatyana Makushok, Jamarc Allen-Henderson, Wallace F. Marshall

## Abstract

Cells have complex and beautiful structures that are important for their function. However, understanding the molecular mechanisms that produce these structures is a challenging problem due to the gap in size scales between molecular interactions and cellular structures. The giant ciliate *Stentor coeruleus* is a unicellular model organism whose large size, reproducible structure, and ability to heal wounds and regenerate have historically allowed the formation of structure in a single cell to be addressed using methods of experimental embryology. Such studies have shown that specific cellular structures, such as the oral apparatus, always form in particular regions of the cell, which raises the question: what is the source of positional information within this organism? By analogy with embryonic development, in which regionalized mRNA is often used to mark position, we asked whether specific regionalized mRNAs might mark position along the anterior-posterior axis of *Stentor*. By physically bisecting cells and conducting bulk RNA sequencing, we were able to identify sets of messages enriched in either the anterior or posterior half and show that RNAi-mediated knockdown of posterior-enriched transcripts corresponding to MYB genes results in a cell’s inability to regenerate the posterior portion of the cell body. We then conducted half-cell RNA-sequencing in paired anteriors and posteriors of cells in which the microtubule cytoskeleton was disrupted by RNAi of β-tubulin or dynein intermediate chains. We found that many messages either lost their regionalized distribution or switched to an opposite distribution, such that anterior-enriched messages in control became posterior-enriched in the RNAi cells, or vice versa. This study indicates that mRNA can be regionalized within a single giant cell and that microtubules may play a role, possibly by serving as tracks for the movement of the messages.

## Introduction

Understanding how complex patterns arise within individual cells is a challenging biological problem due to the vast difference in spatial scales from molecules to whole cells [1,2]. The questions that arise in understanding cellular morphogenesis are essentially the same as those that arise in understanding the development of embryos - how does the system know which structures to make and how to position them relative to an overall body plan? A classic model organism that facilitates the study of developmental biology in a single cell is the giant ciliate *Stentor coeruleus* [3,4]. *Stentor* cells are 1 millimeter long and covered with blue stripes of pigment (Fig 1A) which alternate with longitudinal rows of cilia consisting of basal bodies associated with microtubule bundles (KM-fibers) that run the length of the cell [5]. The cone-shaped cell has an array of cilia known as the membranellar band (MB) at its anterior end and a holdfast at the posterior. Orthogonal to the anterior-posterior (AP) axis defined by these structures, the pigment stripes show a graded distribution of width, defining a circumferential axis. Starting with the widest stripes, the stripe width gradually decreases as one moves counter-clockwise around the circumference, until eventually, the narrowest stripes meet the widest stripes, creating a discontinuity known as the contrast zone.

**Figure 1.**
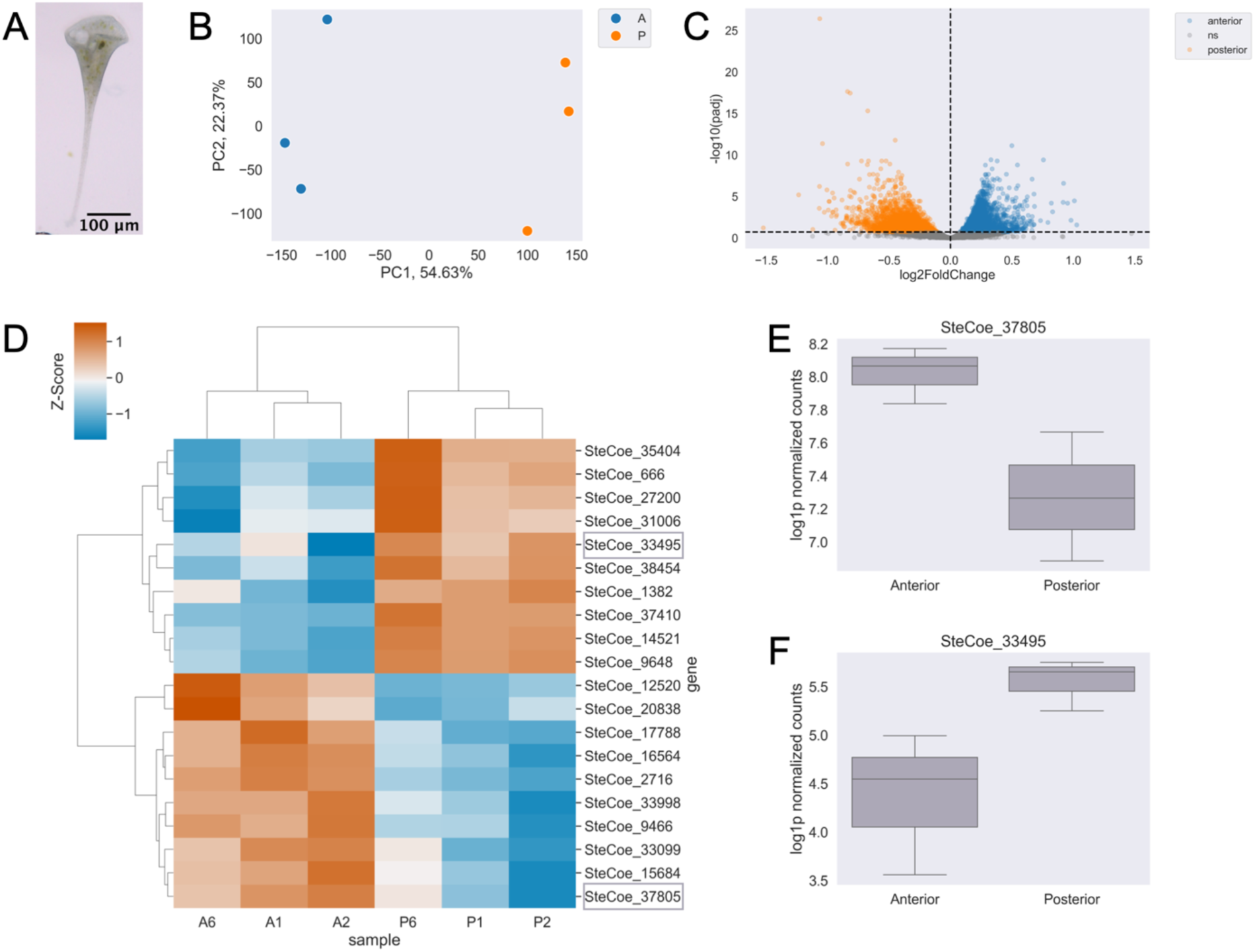
Bulk RNA sequencing reveals RNAs are patterned along the anterior-posterior axis in Stentor coeruleus. (A) Stentor coeruleus. (B) Principal Component Analysis (PCA) showing a clear separation of anterior (color) and posterior (color) samples based on their mRNA abundance profiles. (C) Volcano plot displaying results of differential enrichment analysis of anterior vs. posterior mRNA abundances. Points correspond to individual genes. Statistically significantly differentially enriched genes (q-value < 0.2) in the anterior are blue, posterior in orange, and not significant in gray. (D) Heatmap of relative abundance (z-scores) where rows correspond to individual genes and columns are samples for the top 10 enriched genes in anterior or posterior samples. Red indicates higher enrichment and blue indicates relative depletion. *Representative genes in (E,F) circled in gray. (E) Representative example and boxplot of log-normalized counts of an anterior-enriched message, SteCoe_37805. (F) Representative example and boxplot of log-normalized counts of a posterior-enriched message, SteCoe_33495*.

Part of the reason that *Stentor* was first used as a model system is its ability to regenerate after surgical manipulations. If any part of the *Stentor* cell is cut off, the missing piece regenerates in a matter of hours to yield a normal cell. If a cell is cut in half, each half regenerates its normal structure [6,7]. Part of the reason that *Stentor* can regenerate from these surgical operations is that it contains an elongated macronucleus that contains many copies of every gene. Another reason *Stentor* can regenerate is its ability to employ a range of mechanical wound-closure mechanisms to keep its cytoplasm from leaking out while it patches over a wound [8]. *Stentor’s* huge size, easily visible patterning, and amazing powers of regeneration attracted many developmental biologists during the turn of the last century, including Thomas Hunt Morgan [7]. For almost a century, up until the 1970s, microsurgery was used to analyze morphogenetic processes in *Stentor*, resulting in a wealth of information about how cells respond to geometrical re-arrangements [4,9]. But *Stentor* was never developed as a molecular model system, and we do not currently know the molecular basis for how it can develop or regenerate its structures. To exploit the unique opportunities presented by this organism, we have previously sequenced the *Stentor* genome [10], developed methods for RNAi [11], and analyzed the transcriptional program of regeneration [12]. Proteomic analysis was also reported for isolated MB [13] and transcriptomic analysis was reported for bisected cells of the related species *Stentor polymorphus* [14].

Perhaps the most fundamental question about pattern formation in any organism is what is the source of positional information. That such information exists in *Stentor* can be inferred from the fact that cellular structures always form in defined and reproducible positions [4]. For example, during regeneration or normal cell division, the new MB always forms halfway down the AP axis at the contrast zone between narrow and wide stripes [15]. It is generally true in ciliates that specific surface structures form at positions that can be defined with respect to a cylindrical coordinate system [16–18]. In this system, position is defined by two orthogonal axes - an AP axis that runs along the length of the cell, and a circumferential axis that runs around the equator, which in Stentor is manifest by the gradient of pigment stripe width. We have previously analyzed the distribution of the proteome relative to the AP axis, by cutting individual Stentor cells into pieces and analyzing the fragments by mass spectrometry [19]. This study estimated that approximately 25% of the proteome was polarized relative to this axis.

The role of regionalized mRNA as a source of positional information in embryonic development is well established, particularly in *Drosophila*. We anticipate that regionalized RNAs in different parts of the *Stentor* cell will be important sources of positional information, just like they are in embryos. Unlike the case of *Drosophila* development, where many important genes had already been identified in genetic experiments and served as candidates for localized messages, we do not currently have a list of candidate positional information molecules to map. Inspired by prior work in cryo-slicing *Drosophila* embryos to obtain positional information [20,21], we can use a similar unbiased approach to map the entire transcriptome to identify genes that might have a regionalized distribution.

The unbiased nature of a sequencing-based method for probing localization stands in contrast to microscopy-based methods. Many studies survey message localization using fluorescent *in situ* hybridization (FISH), single-molecule FISH (smFISH [22]) or multiplexed error-robust FISH (merFISH [23]) in cells that lack an obvious polarity axis, so collecting physical fragments of cells representing specific regions and mapping fragments back to any reference frame would not be possible. Consequently, in those cases, each message must be visualized sequentially by imaging, which is time-consuming and expensive. The giant size of *Stentor*, combined with the reproducible cell geometry and easily visualized body axis markers, makes it possible to use a sequencing-based method to map RNA regionalization relative to the body axis coordinates.

The first question is whether genes are regionalized in *Stentor*. If it turns out that RNAs are regionalized to different parts of the *Stentor* cell, the next question is: how might this be achieved? The surface of a *Stentor* is covered with parallel microtubule bundles. Studies in other systems, including yeast and human neurons, show that the cytoskeleton is involved in active RNA transport [24,25]; therefore, we hypothesize that transcripts in this giant cell may be polarized by trafficking specific messages along the cell’s polarized cytoskeletal tracks.

Here, we investigate the regionalization of the *Stentor coeruleus* transcriptome using bulk RNA sequencing of anteriors and posteriors and show that roughly 15% of the transcriptome is polarized along the AP axis. Many of these polarized transcripts encode transcription factors, and we show that knockdown of two MYB transcription factor genes whose transcripts are enriched in the posterior half of intact cells prevents proper posterior regeneration. We also show that knockdown of both β-tubulin and cytoplasmic dynein intermediate chains (henceforth simply referred to as dynein) causes cells to display abnormal cell morphology. To investigate whether disrupting the microtubule cytoskeleton disrupts RNA regionalization, we conducted half-cell RNA sequencing in paired anteriors and posteriors upon knockdown of β-tubulin or dynein. Overall, we find evidence for a global disruption in RNA regionalization upon perturbation of the microtubule cytoskeleton and identify many candidate transcripts with an anti-correlated shift in AP skew.

## Methods

### Stentor culturing

*Stentor coeruleus* were originally obtained from Carolina Biological Supply and have been maintained in the Marshall lab in small Pyrex dishes at room temperature in filter-sterilized pasteurized spring water (Carolina Biological Supply – Burlington, NC), subsequently referred to as <u>C</u>arolina <u>S</u>pring <u>W</u>ater (CSW). Once or twice a week, a lentil-sized pellet of *Chlamydomonas reinhardtii* grown on TAP plates [26] was resuspended in CSW and fed to *Stentor*. Before RNAi, *Stentor* were isolated from the main culture in 6-well tissue culture plates containing 2 mL CSW, starved for 3 days, and washed 3 times with CSW to eliminate residual *Chlamydomonas*.

### RNAi

For sequencing experiments in cells with disrupted microtubules, β-tubulin RNAi was conducted by feeding as previously described [11] with minor changes. RNAi constructs were synthesized by Radiant Inc. using sequences from the *Stentor* genome. The sequence targeting the β-tubulin gene, SteCoe_5719, was amplified with the following oligos: forward - 5’ ATGGTGACTTAAATCACTTGGTTAGTGC 3’, reverse – 5’ TTAAGCAGCTTCCTCTTCCTCATCG 3’. Although *Stentor* has six β-tubulin genes, the high level of sequence conservation is such that this RNAi construct likely targets multiple genes in the family, based on the observation of multiple stretches greater than 20 base identity, which is also supported by our previous demonstration of loss of β-tubulin at the protein level by immunofluorescence [10]. Transformed HT115 *E. coli* were grown to log phase, and expression of dsRNA was induced with 1 mM IPTG overnight at room temperature. Overnight 50 mL cultures were spun down, washed with CSW, and resuspended in 30 mL of CSW. Aliquots of 1 mL were pelleted, with as much supernatant as possible removed, and frozen at -80°C for future use. For feeding, pellets were resuspended in 1 mL CSW. *Stentor* were fed 500 μL control (empty vector) or β-tubulin RNAi bacteria once a day for 7 days.

For sequencing experiments in cells with disrupted dynein motors, dynein intermediate chain RNAi was conducted as above. G04 targets SteCoe_24570 and SteCoe_13560 amplified with forward - 5’ AAGCAGGAACGTGAAGATGC 3’, and 5’ TGCCGATTATTGGGAAAGTG 3’. G05 targets SteCoe_14054 amplified with forward – 5’ AAAACGCTTGCTTGAGCTTC 3’ and reverse – 5’ TGGATGAGTATGGCTTTCTGC 3’.

For experiments to obtain representative images (brightfield and fluorescence) as well as screening Myb DNA-binding domain genes, we used a recently developed method to deliver RNAi by microinjection [27]. This method is more time-efficient and has a higher efficacy than RNAi by feeding, but was not yet established at the time the sequencing experiments were conducted. SteCoe_15300 and SteCoe_33495 constructs corresponding to the first half of each gene were synthesized using Twist Biosciences. Plasmids (2 μg) were linearized with PvuI-HF (NEB), gel-purified with Zymoclean Gel DNA Recovery Kit (Zymo Research), and eluted in 6 μL of water. One ug of linear template was used for *in vitro* transcription with the TranscriptAid T7 High Yield Transcription Kit (Thermo). *Stentor* were incubated in 1.5% methylcellulose for two days before injection. dsRNA at a concentration between 100 and 1000 ng/μL was co-injected into cells with fluorescent dextran using a Nanoject II or Nanoject III (Drummond) and pulled capillary needles as described previously [28].

### Bisection

Bisection is conducted as described previously using pulled capillary needles under a dissection scope for inducing regeneration [29]; however, cells were immediately processed for RNA extraction or sequencing rather than kept for observation.

### Brightfield and Fluorescence Microscopy

For brightfield images, cells were placed in a 13.6 μL droplet onto a slide with a 9 mm diameter x 0.12 mm depth SecureSeal spacer (Grace Bio-Labs) and covered with a coverslip. Images were collected on an AxioZoom V16 (Zeiss) with a 2.3x objective at 32x magnification and a Nikon Rebel T3i SLR Camera. MYB knockdown cells were imaged in low-light settings so as not to disturb the cells and allow them to fully extend. Brightness was adjusted during processing for visualization purposes.

For fluorescence images, MeOH fixed cells were stained with Hoechst 33342 (Thermo), mounted with DABCO in 70% glycerol in PBS, and pipetted onto a coverslip in a 13.6 μL droplet. A slide with a spacer was placed onto the coverslip droplet upside down to ensure the cells remained close to the coverslip for imaging. Images were collected on a Nikon Ti-E inverted microscope with a 25x silicone objective with a 405 nm or 561 nm laser, Hamamatsu Quest camera, and a VT-iSIM super-resolution module. In the images shown, we imaged autofluorescence from the Stentorin pigment, which forms stripes on the cell surface, as a proxy for cortical organization.

### Sequencing

#### Data availability

Raw data will be available on Dryad: https://doi.org/10.6078/D1WT6W

Data were processed and analyzed with custom scripts in Python and R, which are available here: https://github.com/aralbright/2022_AADAWM

#### Bulk RNA sequencing

Within 20 minutes of bisection, 50 anteriors and 50 posteriors were placed in Trizol with as little CSW leftover as possible. RNA was extracted using a Direct-zol RNA Purification Kit (Zymo), and libraries were prepped using the NEBNext Ultra II RNA Library Prep Kit (NEB) following manufacturer’s instructions.

#### Bulk data processing and analysis

We used kallisto [30] to generate a reference index and quantify transcript abundance (Table S1), ComBat [31] to batch correct raw estimated counts (Table S2), and sleuth [32] to perform differential enrichment analysis with a permissive q-value cutoff for significance of 0.2 (Table S3).

#### Half-cell RNA sequencing

Individual *Stentor* that displayed phenotypes were washed and transferred onto parafilm in a 2 μL droplet of CSW and bisected laterally using a needle and processed within 20 minutes of bisection. Cells were lysed and prepped for sequencing with the NEBNext Single Cell/Low Input RNA Library Prep kit for Illumina (Cat. No. E6420) according to the manufacturer’s protocol. Samples were pooled and submitted for sequencing on an Illumina NovaSeq6000 (SP200) or NovaSeqX (SP300). The experiments with β-tubulin and dynein RNAi were done separately, therefore each experiment has its own control samples.

#### Half-cell data processing and analysis

Adapter trimming was performed using flexbar [33] as instructed by NEB for the NEBNext Single Cell Library Prep Kit: https://github.com/nebiolabs/nebnext-single-cell-rna-seq. We used kallisto [30] to generate a reference index and quantify transcript abundance (tubulin – Table S4, dynein – Table S6).

To reduce the effects of outlying transcript abundance on significance, we adjusted log-transformed TPM values by adding the sample mean log-transformed TPM. Anterior-posterior skew is then defined as the difference over the sum of adjusted log1TPM.

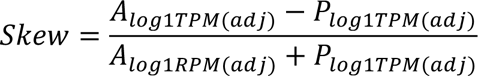

Genes were selected for further analysis that passed a minimal cutoff of abundance, to ensure sufficiently robust statistical comparisons, and it was also required that genes had a coefficient of variation (variance divided by the mean) less than 2 to remove genes with such variable distributions that comparisons might be unreliable. For genes meeting these criteria, skews were compared for both control and β-tubulin RNAi samples, or control and dynein RNAi samples.

We were particularly interested in genes with anti-correlated skews, the most interesting candidates for disrupted regionalization. Such genes were identified with an unbiased feature selection by PCA, in which the genes were treated as factors and the samples as data points, to identify linear combinations of genes that give the highest variation among the samples. Feature selection was then followed by a t-test for significance (tubulin – Table S5, dynein – Table S7).

## Results

### Identification of regionalized messages in Stentor

To ask whether any genes have a regionalized distribution relative to the AP axis, we manually cut *Stentor* cells in half, using the MB as a marker to define the anterior end, and cutting approximately mid-way between the MB and the other end of the cell. For each sample, 50 anteriors and 50 posteriors were processed in bulk RNA-sequencing. The first principal component (PC1) of the principal component analysis (PCA) captures the primary variation in the dataset, with data points corresponding to anterior samples positioned distinctly from posterior samples, suggesting that PC1 effectively represents the AP axis (Fig 1B). Using a permissive q-value threshold of 0.2 for statistical significance and future exploratory analysis of candidate genes, we identify 6292 genes that are statistically significantly differentially enriched between the anterior and posterior half of the cell (Fig 1C, Table S3). This threshold allows for exploratory analysis of more candidate genes in the future, although we acknowledge that a q-value of 0.2 may result in a higher false discovery rate. Our result corresponds to roughly 15% of the total number of gene models.

Log fold change in the positive direction corresponds to messages predominantly enriched in the anterior, with the greatest difference in abundance in SteCoe_37805, SteCoe_33099, and SteCoe_15684. SteCoe_37805 and SteCoe_33099 are 91.2% identical and are predicted to encode membrane transporter proteins. The SteCoe_15684 gene product is a predicted cadherin domain-containing protein. Log fold change in the negative direction corresponds to messages predominantly enriched in the posterior, with the greatest difference in relative abundance in SteCoe_33495, SteCoe_38454, and SteCoe_31006. The protein product of SteCoe_33495 contains Myb-like domains, SteCoe_38454 encodes an FCP1 homology domain-containing protein, and SteCoe_31006 encodes an RGS domain-containing protein. The full list of regionalized genes is given in Table S3.

One hypothesis for why mRNA might be regionalized in different parts of the cell would be to support local translation of proteins needed to build structures in the corresponding region of the cell. We previously reported a proteomic analysis of dissected *Stentor* cells and found sets of proteins that were strongly enriched in either the membranellar band (MB), the anterior of the cell from which the MB had been removed, or in the posterior of the cell [19]. Given the huge size of a Stentor cell, local translation may be necessary to make protein localization more efficient, or else enrichment of transcripts in some region might be a mechanism for producing regional differences in protein composition. In either case, the local translation hypothesis would predict a strong correlation between these regionalized proteins and regionalized mRNAs. However, when we examined genes for which both the message (this study) and protein ([19]) showed a regionalized difference on the AP axis, the majority of regionalized proteins corresponded to anteriorly regionalized messages, even when the proteins were posteriorly regionalized (Table 1). Statistical analysis of this data indicates no significant correlation between message regionalization and protein regionalization (Fishers Exact Test: (A and P) p = 1.0, (A and MB) p = 0.11). Given that the proteomic analysis only identified relatively abundant proteins, one possible explanation might be that the anterior regionalized messages tend to encode abundant proteins. At this point, we do not know how to explain this observed pattern. The first question is whether or not regionalized transcripts play any functional role in generating regionally different patterns within the cell.

**Table 1.** mRNA and Protein Overlap.

|  | **Anterior Protein** | **Posterior Protein** | **Membranellar Band Protein** |
| --- | --- | --- | --- |
| **Anterior RNA** | 23 | 5 | 24 |
| **Posterior RNA** | 1 | 0 | 7 |

### Regionalized transcription factors are necessary for proper cell morphology

Of the many differentially enriched transcripts, 50 correspond to Myb DNA-binding domain proteins, 10 of which are anterior-enriched and 40 of which are posterior-enriched (Table S3). Indeed, one of the most posterior-enriched transcripts discussed above, SteCoe_33495, corresponds to a transcription factor with a Myb DNA-binding domain. Genes in this expanded family of transcription factors are known to control morphogenesis and patterning in plant cells [34] and are well-known proto-oncogenes in humans [35,36]. To ask whether Myb DNA-binding domain transcription factors are important for cellular patterning in *Stentor*, we conducted a targeted RNAi screen of MYB genes found in our study and asked whether these cells regenerated normally.

In each case, following injection of dsRNA, cells were bisected into anterior and posterior half-cells. *Stentor* typically regenerates 8 hours after bisection; however, here we imaged cells at 24 hours post-bisection to ensure they had sufficient time to completely regenerate. For controls we used LF4, a gene that affects ciliary length but not cell morphology, as well as MOB1, which has characteristic effects on cell morphology, to confirm the efficacy of the RNAi procedure. Cells were only counted when the image provided a clear view of the cell in which the MB and posterior were visible. We found that a considerable number of MYB knockdown cells were unable to regenerate a posterior, unlike either of our controls (LF4 and MOB1). As shown in Figure 2, RNAi targeting either Myb-encoding gene causes a fraction of anterior cells to lack the characteristic tapered posterior tail region that gives Stentor its trumpet-like shape. Additional images showing how the shapes are classified in anterior half-cells are present in Supplementary Figure 1. This phenotype was seen in 26% of SteCoe-15300 (6/23) and 40% of SteCoe_22495 (8/20) RNAi anterior half-cells, compared to 10% (2/20) of LF4 and 5% (1/19) of Mob1 anterior halves. In posterior half-cells, this apparent phenotype occurred at a frequency matching what was seen in the controls.

**Figure 2.**
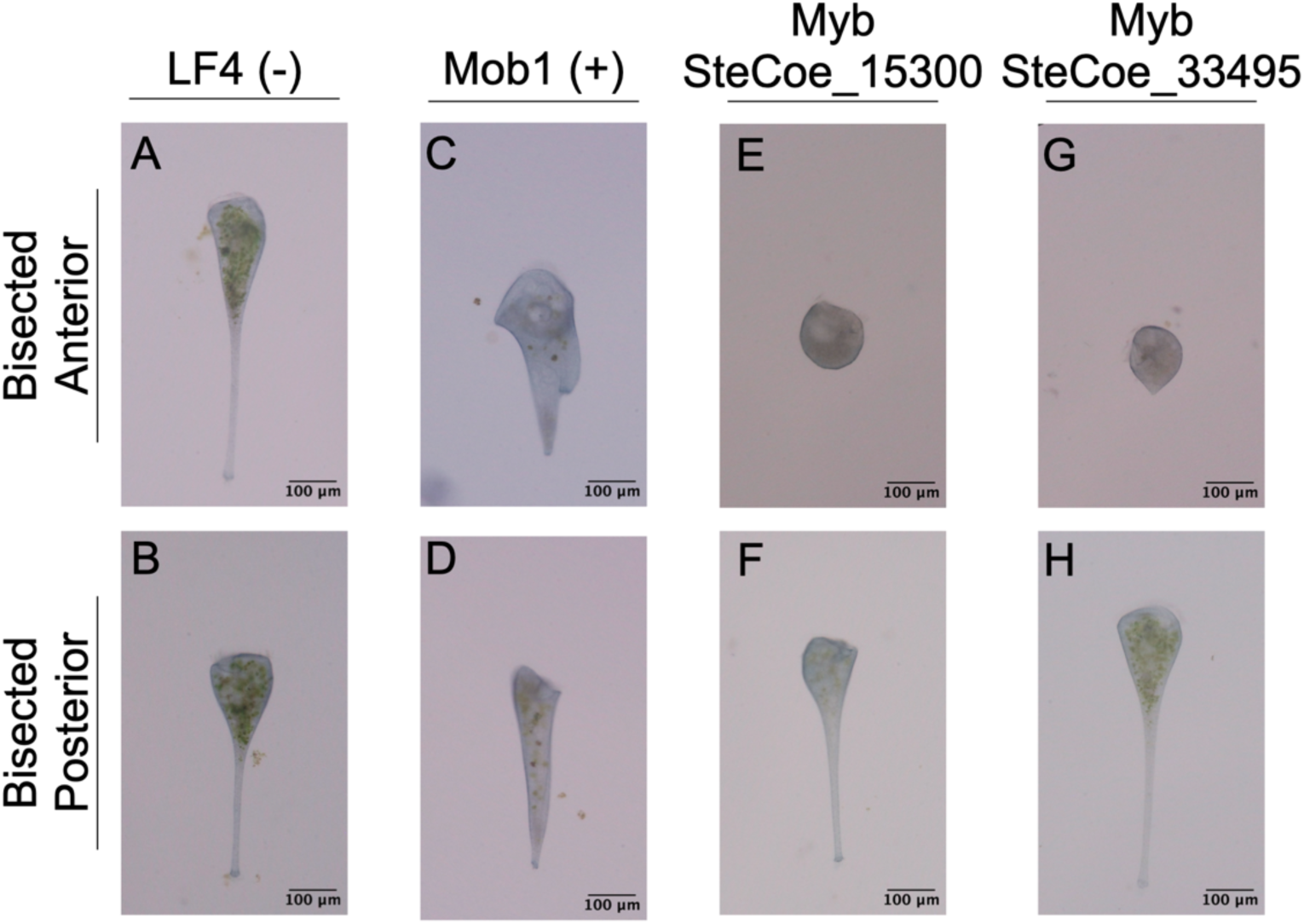
Myb DNA-binding transcription factor knockdown impairs posterior regeneration. Brightfield images of anterior and posterior knockdown cells 24 hours post-bisection. The negative control, LF4 (A - anterior, B - posterior) corresponds to a gene uninvolved in regeneration. The positive RNAi control, Mob1 (C - anterior, D - posterior) corresponds to a gene important for posterior cell morphology but with a distinct phenotype (extra tails and a less tapered body) from what was observed with the Myb constructsSteCoe_15300 (E - anterior, F - posterior) and SteCoe_33945 (G - anterior, H - posterior) correspond to MYB genes.

These results are not consistent with a model in which posterior regionalized transcripts are being stored there for use by posterior half cells when they need to form a new anterior half-cell, but rather suggest that transcripts corresponding to transcription factors may be regionalized in the cell to drive local transcription of genes encoding proteins needed for regenerating structures in that same part of the cell supporting the idea that spatially patterned transcription factors are necessary for proper cellular patterning in *Stentor.* If regionalized transcripts provide part of the explanation for regional differences in protein composition and patterning, this raises the question of how the transcripts themselves become regionalized.

### Role of microtubules in cell morphology

In many cell types, mRNA can be moved along microtubule tracks. If regionalized transcripts are important for cell morphogenesis, we would expect disruption of such tracks to affect cell shape. Prior work has shown that RNAi knockdown of β-tubulin causes cells to lose their typical structure and orderly microtubule bundles [11]. Here, we replicate the findings that knocking down β-tubulin causes cells to lose their characteristic shape and resemble a tulip rather than a trumpet (Fig 3B). In addition, we show that knocking down either a pair of genes encoding a cytoplasmic dynein 1 intermediate chain 2 (SteCoe_24570 and SteCoe_13560) or a gene encoding cytoplasmic dynein 1 intermediate chain 1 (SteCoe_14054) also caused cells to lose their characteristic shape, as well as their ability to extend their holdfast (Fig 3C,D). Using autofluorescence of Stentorin, the pigment that gives *Stentor coeruleus* their blue-green color, we also observe aberrations in structure upon β-tubulin and dynein intermediate chain knockdown. In Fig 3F, we see an underdeveloped MB compared to normal *Stentor*, although the orderly cortical rows appear normal. In Fig 3G-H, we see more extreme defects in the cortical structure upon dynein knockdown.

**Figure 3.**
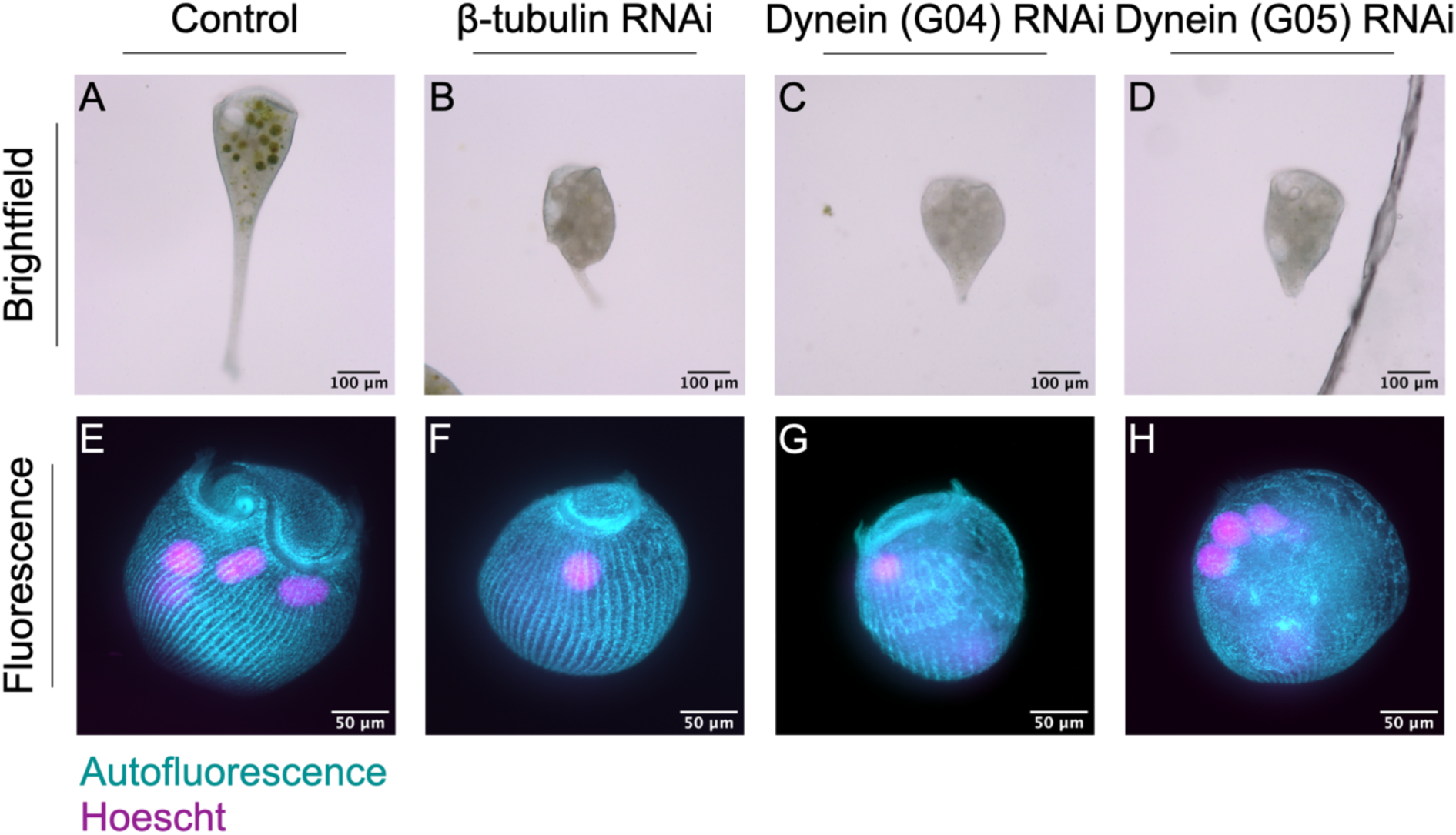
β-tubulin and dynein knockdown results in loss of typical cell morphology. Brightfield images of (A) control, (B) β-tubulin, and (C,D) dynein intermediate chain knockdown cells. Maximum intensity projections for pigment autofluorescence (cyan) and Hoechst staining (magenta) of (E) control, (F) β-tubulin, and (G,H) dynein intermediate chain knockdown cells. Image brightness is adjusted for visualization purposes. G04 and G05 refer to two different RNAi constructs that target distinct dynein intermediate chains encoded by genes SteCoe_24570 and SteCoe_13560 (for G04) and SteCoe_14054 (for G05).

Despite the clear defects in β-tubulin and dynein intermediate chain knockdown cells, the cells remain alive for a period of time and retain a MB at one end and a holdfast at the other. It is thus still possible to recognize the AP axis, allowing us to repeat our analysis of mRNA regionalization by bisecting such cells to produce anterior and posterior half-cells and thereby ask whether disruption of the microtubule tracks or cytoplasmic dynein motors, which we hypothesize to carry messages, has any effect on mRNA regionalization.

### Role of microtubules in mRNA regionalization

To determine whether the microtubule cytoskeleton plays a role in mRNA regionalization, we bisected cells and conducted half-cell RNA sequencing in paired anterior and posterior halves. First, we performed a PCA on control samples to determine if anterior or posterior identity stratifies the samples as shown in Fig 1B for bulk RNA sequencing. in this case, PC2 represents the AP axis (Fig 4A). Next, we projected the β-tubulin knockdown data onto the control PC space (Fig 4B) and found that observable distinction between anterior and posterior samples on PC2 is eliminated upon β-tubulin knockdown (Fig 4C). These differences at the whole-sample level suggest that the microtubule cytoskeleton is broadly important for proper RNA regionalization, such that anterior and posterior samples become more similar to each other in the absence of the microtubule tracks.

**Figure 4.**
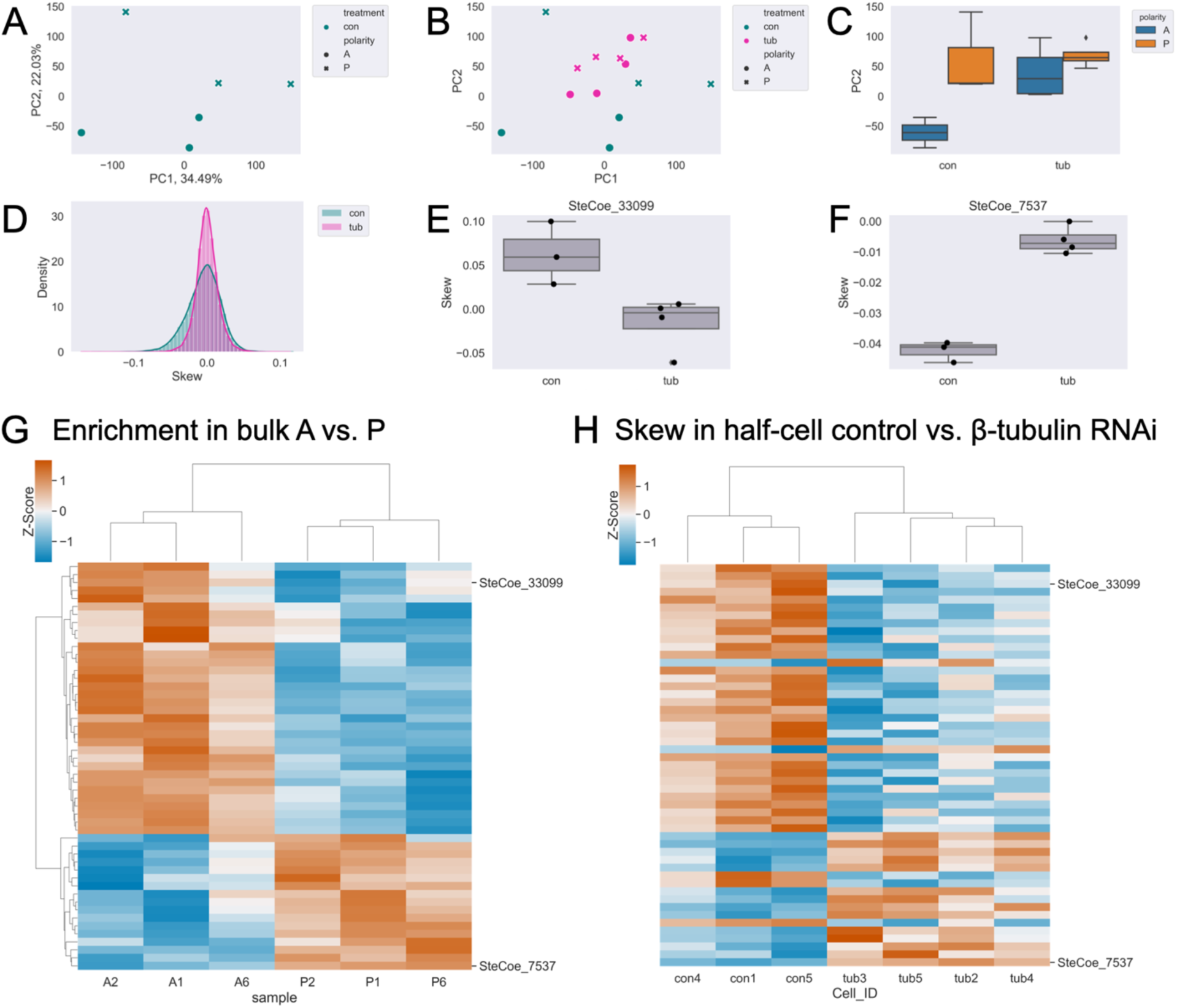
β-tubulin knockdown results in loss of typical cell morphology and mRNA regionalization. (A) PCA on control half-cell RNA data showing a clear separation of anterior (circles) and posterior (x’s) on PC2. (B) Projection of β-tubulin (pink) knockdown half-cells onto the control (green) PC space. (C) Boxplot of PC2 scores for control (left) and β-tubulin (right) data group by anterior (blue) or posterior (orange). (D) Distribution of individual gene skew values in control (green) and β-tubulin RNAi (pink). (E) Representative example, SteCoe_33099, with anterior skew that becomes more posterior upon β-tubulin knockdown. (F) Representative example, SteCoe_7537, with posterior skew that becomes more anterior upon β-tubulin knockdown. (G) Heatmap of relative abundance (z-scores) where rows correspond to individual genes and columns are both differentially enriched in bulk sequencing and *differentially skewed upon on β-tubulin knockdown. Red indicates higher enrichment, and blue indicates lower enrichment. (H) Heatmap of skew (z-scores) in control and β-tubulin RNAi, maintaining the same gene order as (G). Red indicates positive skew corresponding to relative enrichment in the anterior, and blue indicates negative skew corresponding to relative enrichment in the posterior. The four columns on the right cluster the four β-tubulin RNAi cell datasets, and it can be seen that for many genes the skew is now opposite in sign from that seen in controls*.

We then define skew in terms of the difference in abundance of transcripts between the two half-cells in each sample. To quantify polarization along the AP axis, we log(1+x) transformed and adjusted the normalized transcripts per million (TPM) by adding the sample mean log-transformed TPM. Then, we calculated the difference over the sum of AP values for each gene within each sample to obtain a skew value (see Methods for details). We removed genes with highly variable abundance between samples or abundances at the extreme end of high or low to ensure that these characteristics are not affecting our overall calculation of skew. We expect that skews will be variable to some extent even in normally polarized cells, given that the precision of bisection is subject to human error. With this analysis, each gene is assigned an average skew, taken over all samples, with positive values reflecting anterior enrichment and negative values reflecting posterior enrichment. The distribution of skews in β-tubulin knockdown shows a taller, sharper peak, further suggesting that mRNA regionalization is disrupted (Fig 4D).

If microtubule tracks worked to separate a subset of transcripts along the AP axis, one would expect a loss of transcript skew in the RNAi cells, which is certainly seen for many genes. For other genes, in which the skews remain correlated, it suggests that some degree of regionalization is still possible without intact microtubule tracks. The track model can thus account for the loss of correlation or retention of positive correlation. A more surprising outcome would be anti-correlation, such that a gene switches from anterior enriched in the control to posterior enriched in the knockdown, or vice versa. To look for such genes, we performed feature selection by PCA to detect genes with anti-correlated skews (see Methods). Although changes in skew that remain correlated may be significant (i.e. cases where an anterior-enriched gene still has an anterior skew, but perhaps more extreme), candidates with anti-correlated skews upon β-tubulin knockdown are the most interesting for further analysis as these account for the largest differences in skew between control and β-tubulin cells. Following feature selection, we found 156 candidates out of 500 tested with a significant anti-correlated skew (Table S5). SteCoe_33099, a putative membrane transporter and one of the most anterior-enriched mRNAs in bulk sequencing, is skewed more posteriorly upon β-tubulin knockdown (Fig 4E). In the other direction, SteCoe_7537 is a PP2C domain-containing protein that skews more anteriorly upon β-tubulin knockdown. We have previously shown that a PP2A subunit is important for membranellar band, an array of motile cilia at the anterior of the cell, morphogenesis [19], thus SteCoe_7537 may be an interesting gene to examine for its role in posterior patterning.

In addition to SteCoe_33099, we wanted to know if other regionalized mRNAs as determined by bulk RNA sequencing showed an anti-correlated skew upon β-tubulin knockdown. Of the 156 differentially skewed genes upon β-tubulin knockdown, 51 were also differentially enriched in one half of the cell in bulk RNA sequencing (Fig 4G). For most of these genes, including SteCoe_33099 and SteCoe_7537, their skew in control half-cells from the RNAi experiments matches that of the bulk RNA sequencing, but they are oppositely skewed upon β-tubulin knockdown (Fig 4H).

### Role of cytoplasmic dynein in mRNA regionalization

Given that the surface of Stentor is covered with parallel, unidirectional microtubule bundles, an obvious hypothesis is that microtubule motors move transcripts to one end or the other of the cell. There is an extensive literature on cytoplasmic dynein functioning to transport mRNA in other organisms using adaptor proteins like BicD that bind specific messages and couple them to dynein-mediated trafficking [37]. To understand the role of cytoplasmic dynein in cellular patterning, we conducted half-cell RNA-sequencing of paired anteriors and posteriors in control and dynein knockdown *Stentor* using two different dynein RNAi constructs. Construct G04 targets two genes, SteCoe_24570 and SteCoe_13560. G05 targets one gene, SteCoe_14054. In figures and in text, we refer to these as G04 and G05. Similar to in β-tubulin knockdown cells, we see that dynein knockdown cells display altered morphology and disorder in the cortical rows (Fig 3C-D, G-H).

Here again, we performed a PCA on control only samples (Fig 5A), and in this case, PC1 correspond to the AP axis. We then projected the dynein knockdown data onto the control PC space (Fig 5B), and found that observable polarization on PC1 is eliminated upon dynein knockdown (Fig 5C). These results suggest that microtubule motors are also important for proper RNA regionalization on a broad scale. The distribution of skews, specifically with the G04 but not the G05 construct, shows a taller, sharper peak suggesting that perhaps G04 has a greater impact on mRNA regionalization (Fig 5D). Interestingly, we detect 403 anti-correlated skews in G04 (Table S7) following the same method as described above for β-tubulin and none in G05. Whether a lack of significance in G05 is biologically relevant or a result of technical effects is unclear.

**Figure 5.**
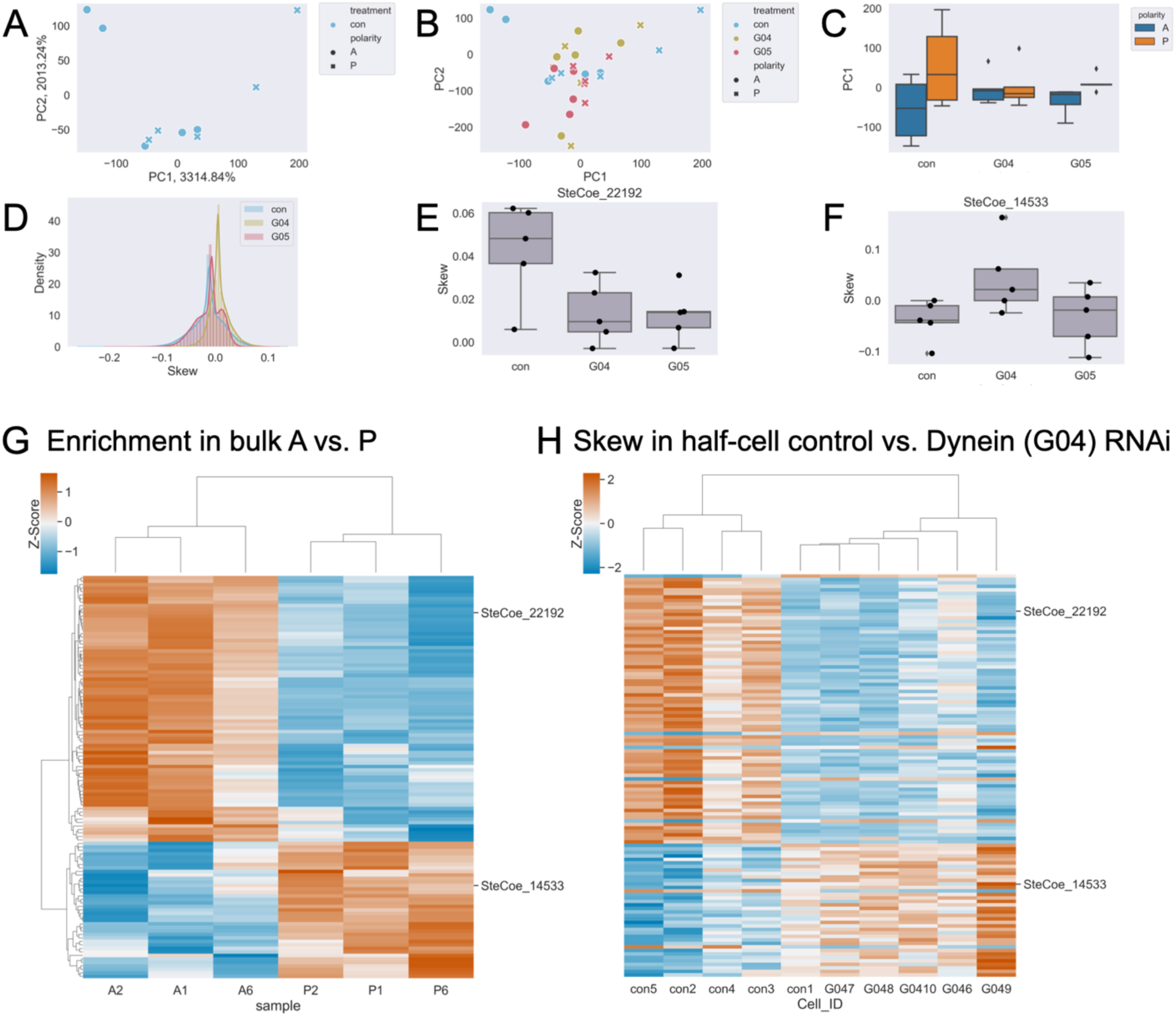
Dynein knockdown results in loss of typical cell morphology and mRNA regionalization. (A) PCA on control half-cell RNA data showing a clear separation of anterior (circles) and posterior (x’s) on PC1. (B) Projection of G04 (yellow) and G05 (red) knockdown half-cells onto the control (blue) PC space. (C) Boxplot of PC1 scores for control (left), G04 (middle), and G05 (right) data grouped by anterior (blue) and posterior (orange). (D) Distribution of individual gene skew values in control (blue), G04 (yellow), and G05 (red). (E) Representative example, SteCoe_22192, with anterior skew that becomes more posterior upon dynein knockdown. (F) Representative example, SteCoe_14533, with posterior skew that becomes more anterior upon dynein knockdown. (G) Heatmap of relative abundance (z-scores) where rows correspond to individual genes and columns are both differentially enriched in bulk sequencing and differentially skewed upon on dynein knockdown. Red indicates higher enrichment and blue indicates lower enrichment. (H) Heatmap of skew (z-scores) in control and dynein RNAi, maintaining the same gene order as (G). Red indicates positive skew corresponding to relative enrichment in the anterior, blue indicates negative skew corresponding to relative enrichment in the posterior. The six columns on the right cluster the 5 dynein RNAi datasets and 1 control cell. Note in this figure that the controls for the dynein RNAi experiment are different from the controls for the β-tubulin RNAi experiment in Figure 3 because controls were performed separately for each RNAi experiment.

SteCoe_22192, one of many genes that encode for actin, mRNA is enriched in the anterior and skews anterior in control half-cell, but is skewed more posterior upon dynein intermediate chain (significant in G04, not significant in G05) knockdown (Fig 5E). SteCoe_14533 is annotated as a protein kinase. Of the 403 differentially skewed genes upon dynein intermediate chain (G04) knockdown, 115 were also differentially enriched in one half of the cell in bulk RNA sequencing (Fig 5G). For most of these genes, including SteCoe_22192 and SteCoe_14533, their skew in control half-cells matches that of the bulk RNA sequencing and they are oppositely skewed upon dynein knockdown (Fig 5H).

## Discussion

Here, we provide evidence that mRNAs are regionalized and the microtubule cytoskeleton is broadly important for proper mRNA regionalization in the giant ciliate, *Stentor coeruleus.* Among all of the regionalized transcripts, we found a large number encoding transcription factors including 50 Myb DNA-binding domain proteins (Table S3). In addition to its role in patterning plant cells and as a proto-oncogene in humans, MYB is also a key regulator of ciliogenesis and centriole amplification during the formation of multiciliated epithelia [38,39], both of which take place during morphogenesis and regeneration in *Stentor.* Upon knocking down two Myb genes, SteCoe_15300 and SteCoe_33495, we found that a considerable number of cells are unable to regenerate their posterior. It may seem surprising to find such messages that encode transcription factors destined to enter the nucleus, regionalized within a cell that has a single nucleus. However, we note that although *Stentor coeruleus* has a single macronucleus, this macronucleus is highly polyploid and spatially extended, running the length of the whole cell. Moreover, the macronucleus takes the shape of a set of nodes joined by small junctions, like beads on a string, such that different nodes might be able to sustain different programs of gene expression. We speculate that transcription factors encoded by regionalized messages might function to locally drive patterns of gene expression in different parts of the macronucleus.

Knockdown of β-tubulin, a major component of the microtubule cytoskeleton, disrupts normal cell morphology and causes cells to lose their orderly cortical rows (Fig 3, [11]). Half-cell RNA-sequencing in paired anteriors and posteriors of the cell also reveals global disruptions in regionalization by PCA upon knockdown of β-tubulin or dynein intermediate chains (Fig 4A-C, Fig 5A-C). Overall, we find many candidates with significant and anti-correlated skews (β-tubulin – Table S5, dynein (G04) – Table S7). However, despite an apparent loss of regionalization in one of our dynein intermediate chain knockdowns globally (G05 – Fig 5C), we find no statistically significantly different transcript skews.

We speculate that transcripts with significant and anti-correlated skews upon β-tubulin or dynein knockdown (G04) may be important for cellular patterning, given the morphological defects and change in transcript skew upon knockdown, and the connection between patterned mRNAs and cell patterning in other contexts. We do note, however, that for a small number of genes, the skew in the control half-cells was opposite to that seen in bulk RNA sequencing, but then became the same as in bulk RNA sequencing in the RNAi cells.

This work shows that mRNA regionalization is disrupted upon the loss of β-tubulin or dynein intermediate chains, but raises several additional questions: (1) Are the messages we found directly involved in cellular patterning or the regeneration of new patterned structures? (2) Is mRNA regionalization by the microtubule cytoskeleton essential for regeneration, given its role in patterning the cell? (3) Does protein enrichment in one half over the other arise from differential transcript regionalization or possibly translation? An analysis of the data in Table 1 showed a lack of correlation between protein regionalization and transcript regionalization, which argues against this idea.

RNA regionalization could be a key to understanding pattern formation and regeneration in *Stentor*. Our results suggest a potential model for how this regionalization is achieved. Given that RNAi knockdown of posterior-regionalized transcripts corresponding to transcription factors inhibits proper posterior regeneration and that disruption of the cortical microtubule bundles led to a loss or reversal of polarized regionalization for many messages, we propose that messages may both be transcribed near where they are needed in the cell and associate with microtubule motors via RNA-binding adaptors, such as BicD, and then move to the ends of the bundles (Figure 6). Future work will be needed to test this model by interfering with kinesin as well as potential adaptor proteins.

**Figure 6.**
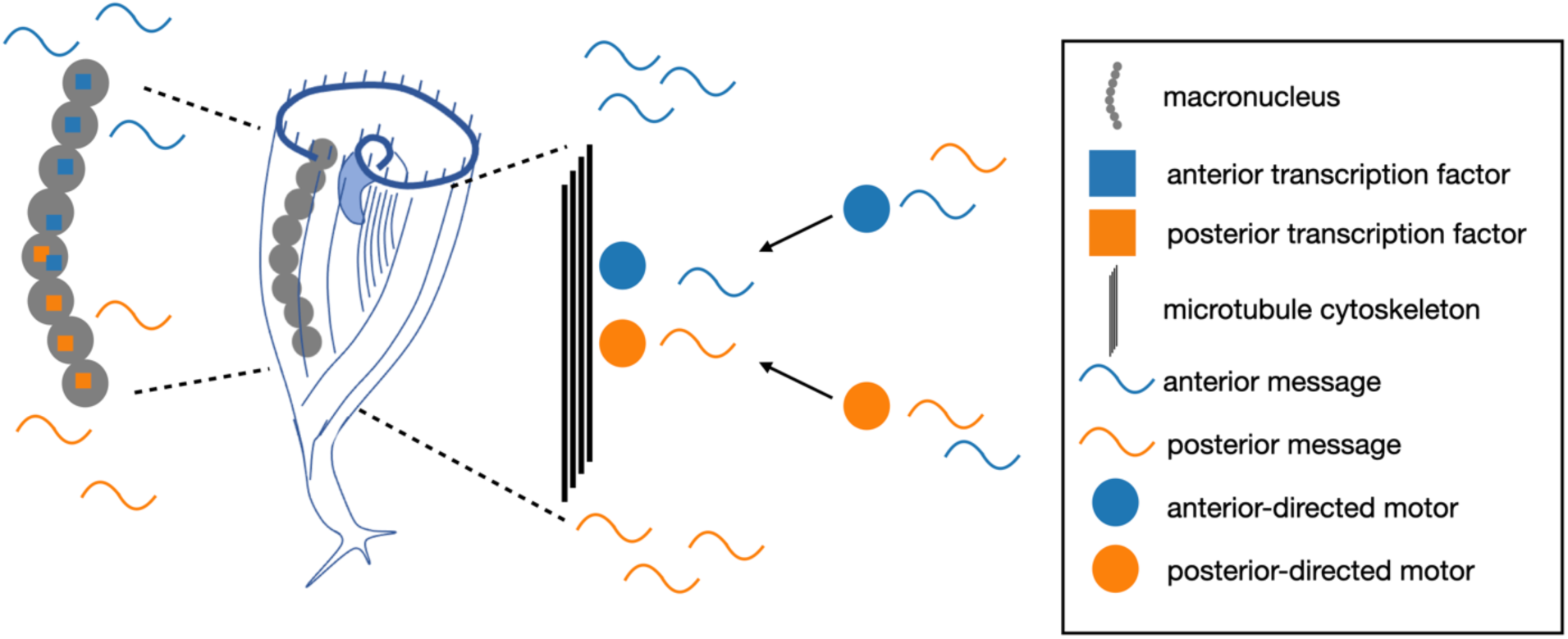
Model for RNA regionalization in Stentor via regionalized transcription factors and/or motor-directed transport.

As this is a preprint-in-progress, we welcome input on additional interesting candidates for further analysis, as well as any feedback or other suggestions for this manuscript.

## Acknowledgements

We would like to acknowledge Mark Slabodnick, Dept. Biology Knox College for his pioneering work on *Stentor* RNAi by both feeding and microinjection. We thank Dyche Mullins, Tanner Fadero, and Sam Lord for microscopy assistance and use of the iSIM. We would also like to thank members of the Marshall Lab, Mark Slabodnick, and Xin Xiang for helpful discussion and feedback. Sequencing was performed at the UCSF CAT, supported by UCSF PBBR, RRP IMIA, and NIH 1S10OD028511-01 grants. A.A. was supported by NIH 5K12GM081266 and NIH K99GM154059. C.Y. was supported by an NSF-GRFP fellowship. This work was supported by NIH R35 GM130327, on which W.F.M. is the principal investigator.

## Note

This is a preprint and is not peer reviewed

## Supplementary Files

Supplementary Table 1. Kallisto raw data table – bulk AP RNA sequencing

Supplementary Table 2. Batch corrected counts – bulk AP RNA sequencing

Supplementary Table 3. Differential enrichment analysis with PyDeSeq2 – bulk AP RNA sequencing

Supplementary Table 4. Kallisto log normalized tpm table – tubulin half-cell RNA sequencing

Supplementary Table 5. Skew analysis – tubulin half-cell RNA sequencing

Supplementary Table 6. Kallisto log normalized tpm table – dynein half-cell RNA sequencing

Supplementary Table 7. Skew analysis – dynein (G04) half-cell RNA sequencing

## Supplementary Figures

**Supplementary Figure 1.**
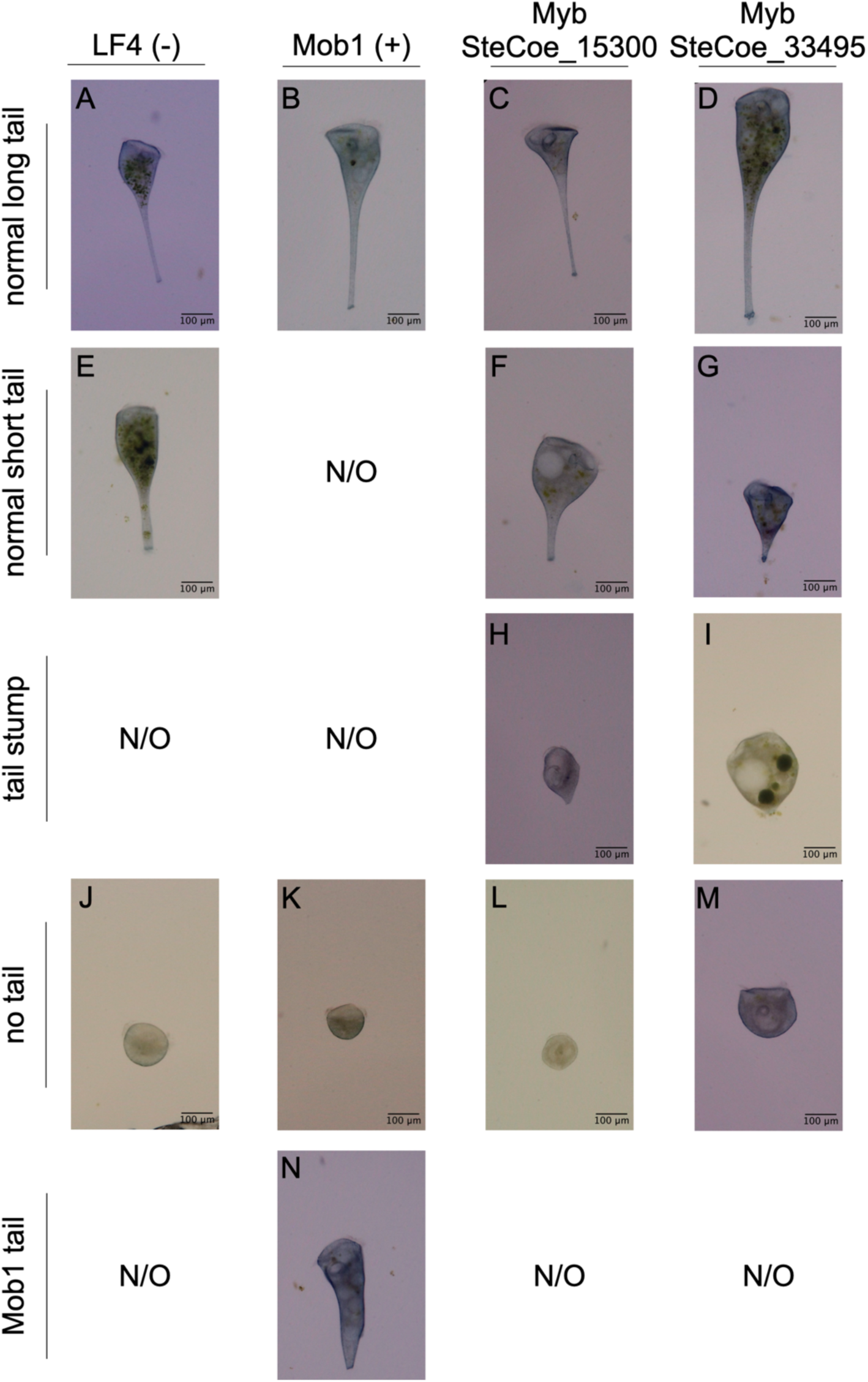
Representative phenotypes from control and RNAi knockdown conditions in anterior half-cells. Brightfield images of anterior halves of knockdown cells 24 hours post-bisection. Normal long tails with tapered shape and *holdfast appeared in all conditions (A-D). Normal short tails with a tapered shape and holdfast appeared in (E) LF4, (F) SteCoe_15300, and (G) SteCoe_33495. Appearance of a posterior stump with a holdfast but no tapered region, which was counted as no tail in our analysis, only appeared in MYB knockdown cells (H) SteCoe_15300 and (I) SteCoe_33495. Cells with no visible tail were occurred in all conditions at least once (J-M). The characteristic Mob1 phenotype characterized by a thick cylindrical body shape was only seen in (N) Mob1 knockdown cells. N/O = not observed*.

